# Carbon Burial in Sediments below Seaweed Farms

**DOI:** 10.1101/2023.01.02.522332

**Authors:** Carlos M. Duarte, Antonio Delgado-Huertas, Elisa Marti, Beat Gasser, Isidro San Martin, Alexandra Cousteau, Fritz Neumeyer, Megan Reilly-Cayten, Joshua Boyce, Tomohiro Kuwae, Masakazu Hori, Toshihiro Miyajima, Nichole N. Price, Suzanne Arnold, Aurora M. Ricart, Simon Davis, Noumie Surugau, Al-Jeria Abdul, Jiaping Wu, Xi Xiao, Ik Kyo Chung, Chang Geun Choi, Calvyn F.A. Sondak, Hatim Albasri, Dorte Krause-Jensen, Annette Bruhn, Teis Boderskov, Kasper Hancke, Jon Funderud, Ana R. Borrero-Santiago, Fred Pascal, Paul Joanne, Lanto Ranivoarivelo, William T. Collins, Jennifer Clark, Juan Fermin Gutierrez, Ricardo Riquelme, Marcela Avila, Peter I. Macreadie, Pere Masque

## Abstract

The hypothesis that seaweed farming contributes to carbon burial below the farms was tested by quantifying burial rates in 20 seaweed farms distributed globally, ranging from 2 to 300 years in operation and from 1 ha to 15,000 ha in size. This involved combining analyses of organic carbon density with sediment accumulation rate in sediments below seaweed farms relative to reference sediments beyond the farm and/or prior to the farm operation. One in every four farms sampled was set over environments that export, rather than retain materials. For the farms that were placed over depositional environments, where sediment accumulation could be quantified, the thickness of sediment layers and stocks of carbon accumulated below the farms increased with farm age, reaching 140 ton C ha^-1^ for the oldest farm, and tended to exceed those in reference sediments beyond the farm and/or prior to the operation of the farms. Organic carbon burial rates in the farm sediments averaged (± SE) 1.87 ± 0.73 ton CO_2_ equivalent (CO_2-eq_) ha^-1^ year^-1^ (median 0.83, range 0.10 – 8.99 ton CO_2-eq_ ha^-1^ year^-1^), twice the average (± SE) burial rate in reference sediments (0.90 ± 0.27, median 0.64, range 0.10-3.00 ton CO_2-eq_ ha^-1^ year^-1^), so that the excess organic carbon burial attributable to the seaweed farms averaged 1.06 ± 0.74 ton CO_2-eq_ ha^-1^ year^-1^ (median 0.09, range −0.13-8.10 ton CO_2-eq_ ha^-1^ year^-1^). This first direct quantification of carbon burial in sediments below seaweed farms confirms that, when placed over depositional environments, seaweed farming tend to sequester carbon in the underlying sediments, but do so at widely variable rates, increasing with farm yield.

## Introduction

Seaweed farming, growing at about 7.3 % year^-1^, accounts for about half of total mariculture production (FAO 2022) and provides a novel opportunity to advance multiple sustainable development goals (Duarte et al. 2022a). These include climate action (Duarte et al. 2022a), with seaweed farming considered an emerging Blue Carbon option (Lovelock and Duarte 2019). Blue carbon, the strategy to mitigate climate change through the conservation and restoration of vegetated coastal habitats (Duarte et al. 2013), is a growing component of climate action delivered by projects targeting mangroves, seagrass and saltmarshes (Macreadie et al. 2021). The climate mitigation capacity from protecting and restoring those blue carbon habitats is estimated at ~3% of the removal of global green-house gas emissions required to achieve the goals of the Paris Agreement (Macreadie et al. 2021). However, seaweed farming can be conducted sustainably over expansive areas estimated to be as large as 677.832 Km^2^ by 2050 (Duarte et al. 2022a). If achieving this scale, seaweed farming could deliver the removal of an estimated 0.24 Pg CO_2_-equivalent (CO_2-eq_) per year stored in sediments below the farms (Duarte et al. 2022a). However, these estimates are based on indirect assumptions as there is, to-date, not a single published empirical estimate of carbon burial in sediments below seaweed farms, rendering the estimates of the potential contribution of seaweed farming hypothetical.

Wild algal forests extend over 7.2 million Km^2^ and have a global production of about 1.32 Pg C year^-1^ (Duarte et al. 2022b), sequestering an estimated 634 Tg CO_2-eq_ year^-1^, which is as much as mangroves, seagrass and saltmarshes combined (Krause-Jensen and Duarte 2016). Seaweeds release a significant fraction of their net primary production (NPP) as both dissolved and particulate organic carbon, exporting, on average, about 43 % of their NPP (Duarte and Cebrian 1996). However, as most algal forests grow on exposed rocky shores, the production exported is advected far away, with seaweed detected in the open sea and deep sea (Krause-Jensen and Duarte 2016, Ortega et al. 2019, Hurd et al. 2022). In contrast, seaweed farms are often located in sheltered bays suspended over soft sediments, which preclude damages due to storms and swells while providing a potential depositional environment where carbon detached from the seaweed can be buried in sediments below the farm. Assuming seaweed farms to behave as wild seaweed populations, Duarte et al. (2022) calculated that seaweed farms could sequester about 11 % of their net primary production (Krause-Jensen and Duarte 2016) or about 3.5 ton CO_2_ ha^-1^ year^-1^, assuming relatively high farm yields. However, this potential has not yet been quantified.

Here we provide a first assessment of organic carbon burial in sediments below seaweed farms. We do so on the basis of a global study including 20 seaweed farms, none of them designed to sequester carbon, located in 11 nations across all continents. Our study supports the hypothesis that most seaweed farms bury carbon in the underlying sediments and that sustainable seaweed farming may contribute to climate change mitigation as a co-benefit to their primary role in helping alleviate hunger and poverty and meet several other sustainable development goals (Duarte et al. 2022a). Carbon burial in sediments below seaweed farms ranged widely, and showed a tendency to be higher when farms placed in physiographic settings conducive to retaining materials in sediments, best reflected in organic-rich muddy to fine sandy sediments, were exploited at high seaweed yields. The high variability in carbon burial rates among farms identifies opportunities to maximize carbon burial by placing seaweed farms in suitable depositional environments. For instance, many farms in Japan are placed over rocky shores, where carbon is likely exported, but not sequestered in sediments below the farm.

The farms in our study (Fig. 1) ranged widely in size, from very small farms with one or a few hectares farmed in Western nations to a 150 km^2^ farm in Ningde, China, which can be seen from space (Tables 1 and 2). The farms ranged in age from just a few years to 320 years of operation in Tokyo Bay, Japan (Tables 1 and 2). Indeed, the farms in the Europe and North America were all very recent and had a small footprint, attesting to the incipient nature of seaweed farming in most Western nations (Tables 1 and 2). The studied farms included 12 of the most important farmed seaweed species, ranging from large, slow growing kelps (*Saccharina latissima*) to fast-growing sea lettuce (*Ulva spp*.). Current yields ranged widely, from 1 to 150 tons fresh weight ha^-1^ year^-1^ (Tables 1 and 2), reflecting the number of crops and harvests per year, ranging from 1 in high-latitude farms to 6 per year in Indonesia, as well as the convenience for the farmers. Concessions or leases are often much larger than the cropped area, and farmers usually set generous spacing among the seaweed lines to prevent the spread of disease, maximize seaweed growth and ease navigation with their vessels, but resulting in reduced seaweed yield per hectare.

**Fig. 1.**
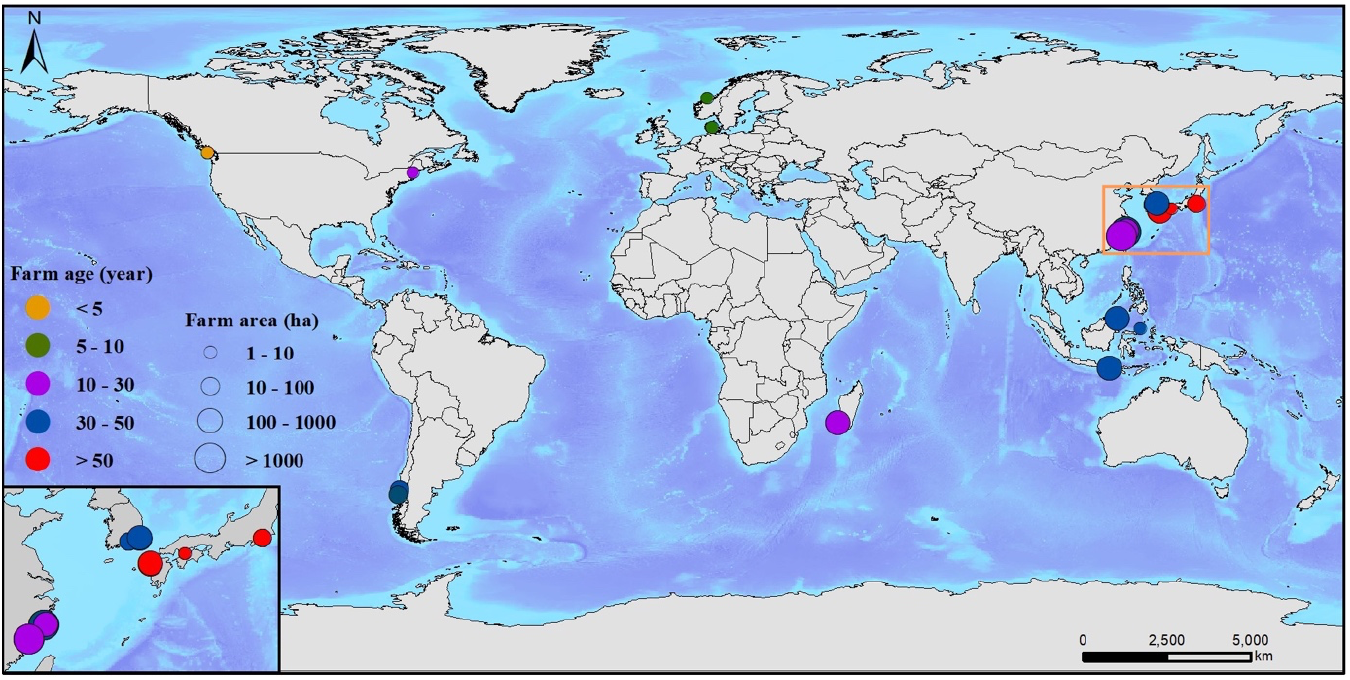
Location of the sampled seaweed farms, the time (years) elapsed since the onset of farming and the size of the harvested area (ha) (Tables 1 and 2). The orange square in Asia shows the area highlighted in the insert.

**Table 1.**
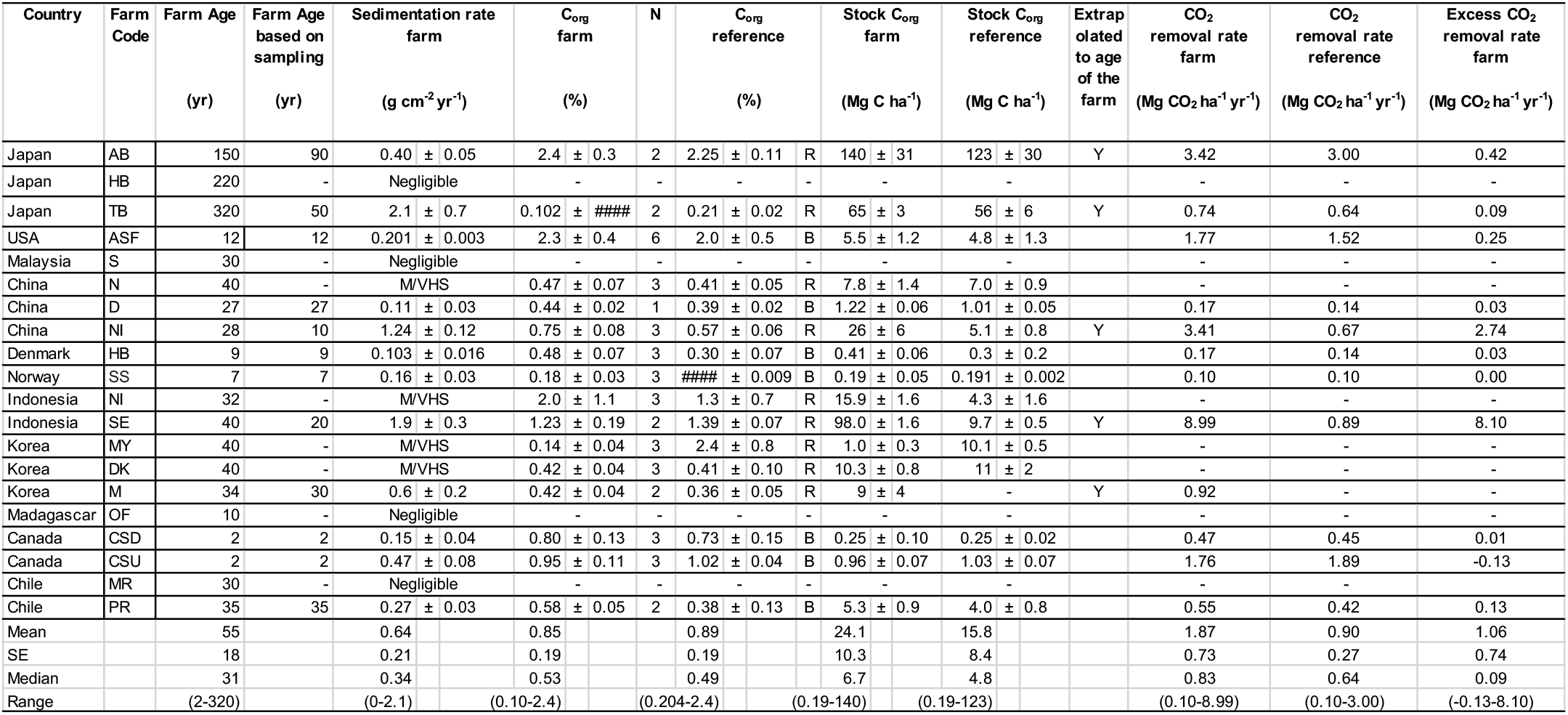
Time (years) since the onset of operation of the seaweed farms and as recorded with the sediment cores based on ^210^Pb dating, sedimentation rates in the farm (mean ± SE), average sediment organic carbon concentration (C_org_) in sediments deposited during the farm operation or in reference sediments, organic carbon stock accumulated in sediments below the farm during the farm operation and CO_2_ equivalent burial rate in sediments deposited during the farm operation or in reference sediments, along with the mean, standard error (SE), median and range of values across the sample set. N = number of cores used in estimates. The C_org_ contents, stocks and removal rates in reference were calculated from measurements in cores taken away from the farm (R) or from measurements in cores below the farm at depths before to the onset of farming activities (B). Negligible indicates that sedimentation rates were too low to be resolved, while M/VHS indicates that sediments were either mixed or sedimentation rates were too high to be resolved with the length of sediment core collected.

**Table 2.**
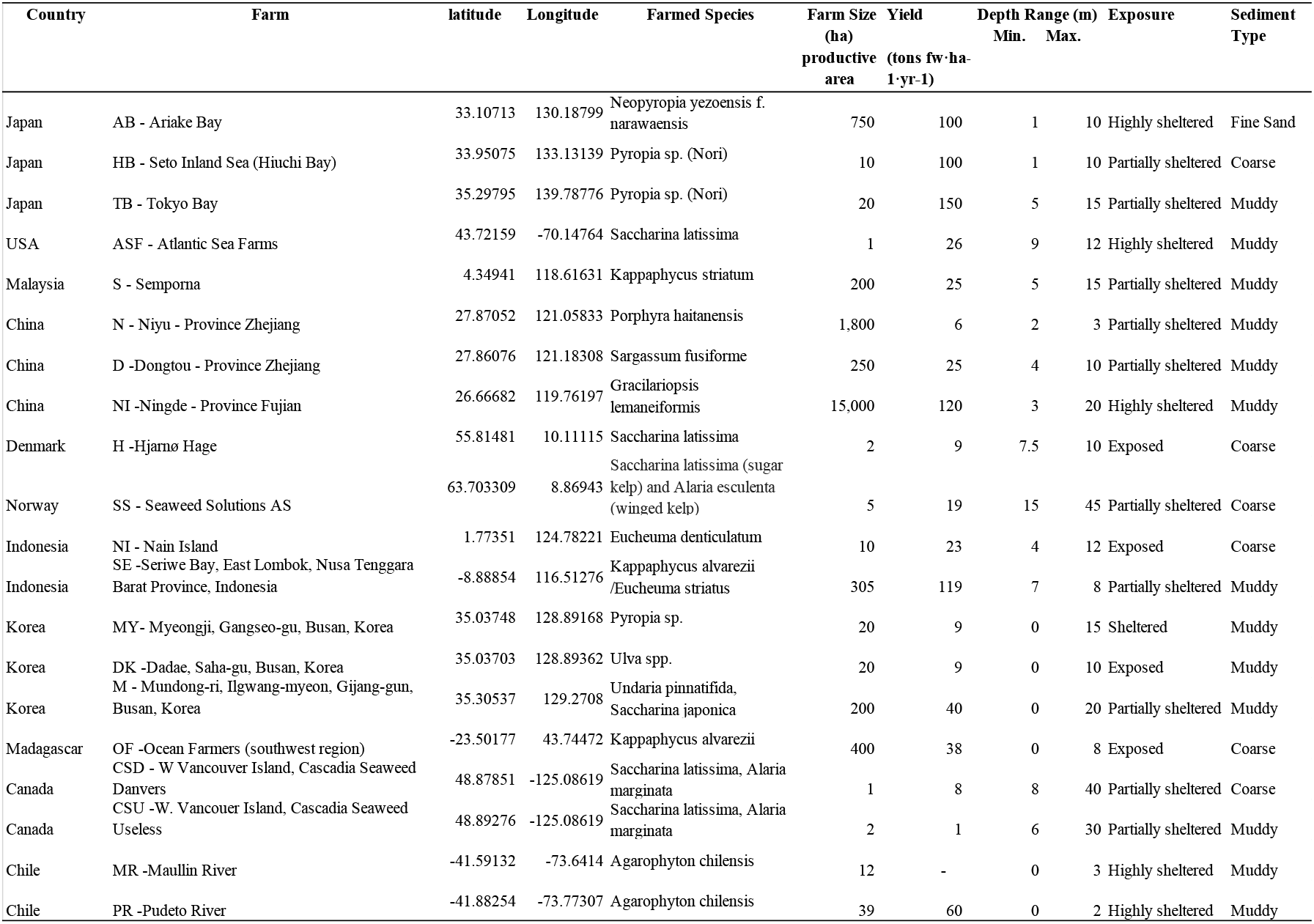
Location (latitude and longitude in decimal degree units), depth range (minimum and maximum) of the farmed area and the current farmed species and the size and yield, as well as exposure and sediment type of the seaweed farm.

The settings of the farms also varied widely, from sites with strong currents and exposure to waves, such as those in Norway, Korea and Indonesia, which maximize nutrient supply and, therefore, seaweed growth, but export, rather than accumulate, materials, to sheltered bays in Canada. Depth also varied from intertidal farms in China and Indonesia to farms located over 30 m of water in Norway and Canada (Tables 1 and 2). Accordingly, the sediments below the farms ranged from muddy to gravel, with coarse sediments indicative of environments exporting rather than retaining materials, including seaweed fragments. Indeed, exposed sites tended to have coarse sediments, compared with sheltered sites, which tended to have muddy sediments, with fine sands dominating partially sheltered sites (χ^2^, P=0.008).

The distance between the center of farms and the reference sites outside the farm was considerable for the larger farms, which occupied all the suitable environment for seaweed farming. As a result, areas away from the farms often differed in physical conditions, reflected in differences in the characteristics of sediments compared to those below the farms. Hence, sediments away from the farm, did not always provide an appropriate reference to compare the effect of seaweed farming on organic carbon stocks and burial rates. In cases where differences in sediment properties prevailed, and the suggested reference sediments could not be compared to sediments below the farm, we compared the upper part of the sediment cores corresponding to sediment accumulated since the farm started operating and the lower part corresponding to sediment accumulated before operations, where the sediment cores reached to this layer.

Sedimentation rates could be resolved based on ^210^Pb concentration profiles in 11 of the 20 farms sampled. Five farms showed negligible sedimentation rates, suggesting they were set in environments that export rather than accumulate materials (Table 1). Four farms had ^210^Pb concentration profiles that denoted either mixing, likely due to bioturbation, or sedimentation rates too high to be resolved with the sediment core length collected (Arias-Ortiz et al. 2018) (Table 1). The estimated thickness of the sediment layer deposited during the operation of the farm ranged widely and increased with farm age (Fig. 2a) for those farms placed within depositional environments. Hence, the sediment layer deposited during farm operations was very thin (0 – 1.1 cm) in farms in Europe and North America, where seaweed farming is a recent development. In contrast, we estimated the sediment layer deposited during farm operations to be about 112 cm thick in the farm in Ariake Bay, Japan, the oldest (150 years of operation) farm located in a depositional environment (Tables 1 and 2).

**Figure 2.**
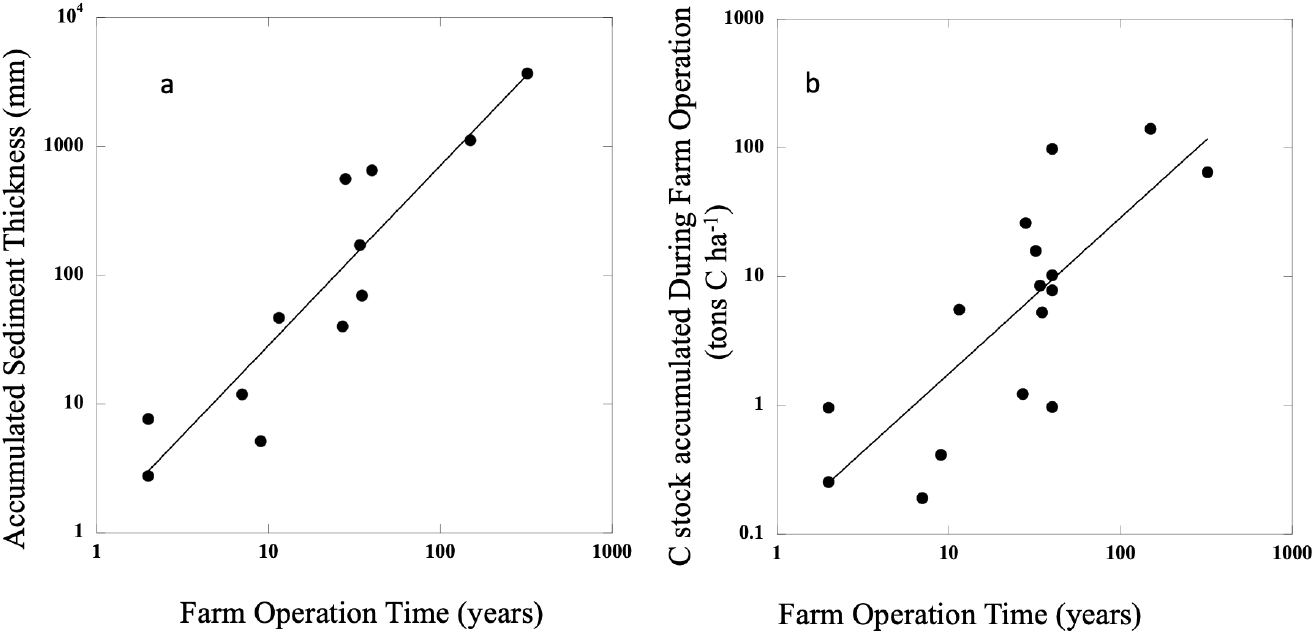
The relationship between the time elapsed since seaweed farming operations were initiated and (a) the thickness of the sediment layer deposited when detectable, and (b) the corresponding stock of organic carbon accumulated per unit farm area. The thickness and carbon stock deposited during farm operations were extrapolated linearly from the sediment layer analyzed (Table 2) when this did not include the entire operation period of the farm. The solid lines represent the fitted regression lines: (a) log_10_ Accumulated Sediment Thickness (mm) = 0.06 + 1.39 (± 0.19) log_10_ Farm Age (years) (R^2^=0.84, P < 0.0001); (b) log_10_ C accumulated during farm operation (tons C ha^-1^) = −0.96 + 1.21 (± 0.16) log_10_ Farm Age (years) (R^2^ = 0.61, P = 0.0004).

Sediment organic carbon concentration averaged (±SE) 0.85 ± 0.19 % of dry weight across farms (median 0.53, range 0.10-2.40%, Table 1). Two-thirds of the farms contained elevated organic carbon concentrations in sediments below the farm corresponding to the period of operation relative to reference sediments beyond the farm or sediment layers deposited prior to the operation of the farms (Wilcoxon ranked paired test, P <0.05, Table 1). The stock of organic carbon accumulated during farm operations in farms that were set over depositional environments ranged from 0.19 to 140 ton C ha^-1^, increasing with the timespan the farms had been in operations (Fig. 2b, Table 1), and tended to exceed the stock of carbon accumulated over the same time in reference sites away from the farm or prior to farm operations (Wilcoxon ranked paired test, P <0.005, Table 1).

Organic carbon burial rates in the farm sediments averaged (± SE) 1.87 ± 0.73 ton CO_2-eq_ ha^-1^ year^-1^ (median 0.83 ton CO_2-eq_ ha^-1^ year^-1^, range 0.10 – 8.99 ton CO_2-equiv_ ha^-1^ year^-1^), and showed a weak tendency to increase with increasing current farm yield (Fig. 3a) and exceed organic carbon burial rates in reference sites away from the farm or prior to farm operations (Wilcoxon ranked paired test, P <0.005, Fig. 3b). The excess organic carbon burial relative to reference sediments, attributable to the operation of the seaweed farm, averaged (± SE) 1.06 ± 0.74 ton CO_2-eq_ ha^-1^ year^-1^ (median 0.09 ton CO_2-eq_ ha^-1^ year^-1^, range −0.13 – 8.10 ton CO_2-eq_ ha^-1^ year^-1^).

**Figure. 3.**
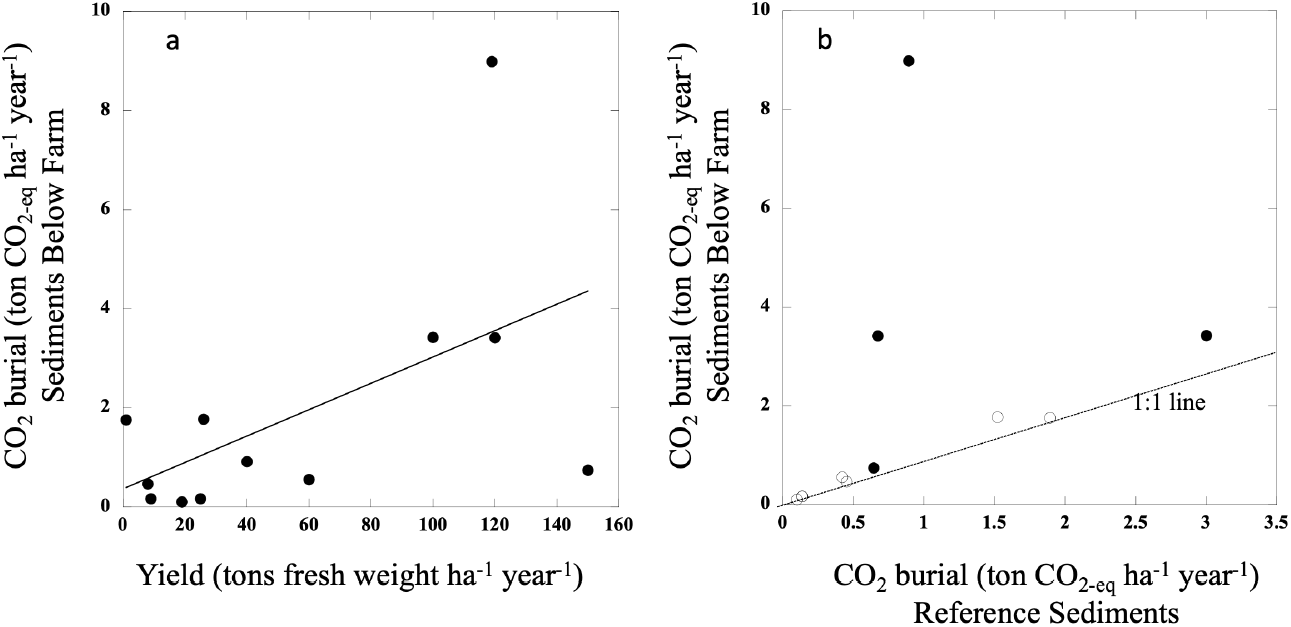
The relationship between the CO_2-eq_ burial in sediments below the farm and (a) the current farm yield and (b) the CO_2-eq_ burial in reference sediments. The solid line in (a) represents the fitted regression line: (a) CO_2_ burial (tons CO_2-eq_ ha^-1^ year^-1^) = 0.37 + 0.026 (± 0.012) Yield (tons fresh weight ha^-1^ year^-1^) (R^2^ = 0.30, P = 0.065). The dotted line in (b) shows the 1:1 line. The empty symbols in (b) denote farms where reference values were obtained from analyses of sediment layers before farm operation.

The results presented are the first direct, empirical estimates of carbon burial in sediments attributable to the operation of seaweed farms. Seaweed farming is a recent development in Europe and North America, where seaweed farms are small and young. This is in contrast to a centenary record of operation in Asia, particularly in Japan, where the farm in Tokyo Bay, in operation for 320 years, had accumulated, extrapolated from the sedimentation rates during the operation of the farm, an impressive 369 cm layer of sediment containing 56 ton C ha^-1^ The average rate of carbon burial in sediments below seaweed farms reported here is within the lower range of the amount of carbon that have been modelled to deposit below seaweed farms (in Norway, Broch et al. 2022), and within the range of what can be expected based on the carbon export from seaweed farms (Fieler et al. 2021, Zhang et al. 2012) and an approximately 10% burial efficiency (i.e. percentage of carbon deposited in the seafloor that is buried in sediments, Middelburg 2019). The estimated carbon burial due to seaweed farming provides a conservative assessment of the role of seaweed farming in carbon sequestration, as particulate organic carbon detached from seaweeds may be outwelled beyond the footprint of the farm by advective and turbulent mixing, as shown for seagrass meadows (Kennedy et al. 2010) and mangroves (Dittmar et al. 2006), and because the estimate does not consider dissolved organic carbon exported to the deep sea, which can be an important component of seaweed carbon sequestration (Krause-Jensen and Duarte 2016). Further, this average carbon burial below seaweed farms corresponds to about 39% to 33% of the estimated average carbon sequestration rate in seagrass and mangrove soils, respectively (Duarte et al. 2013). Applying the average carbon burial below seaweed farms found in our study to the estimated present global area of seaweed farms of 1,983 km^2^ in 2020 (Duarte et al. 2022a) implies that soils below seaweed farms may currently sequester about 0.4 million tons of CO_2-eq_ annually, which could grow, following current practice, to 140 million tons of CO_2-eq_ annually if seaweed farming expanded at the upper growth rate to reach 677.832 km^2^ by 2050 (Duarte et al. 2022a). These estimates need be corrected for the background carbon burial in reference sediments to assess the carbon removal “additionality” contributed by the seaweed farms, which represents about half of the total carbon burial below seaweed farms (Table 1). The estimate of the excess organic carbon buried in sediments below farms reported here represents a crude approximation of the “additionality” to carbon burial contributed by seaweed farms. This is because the use of reference sediments, whether away or before the farm operations started, involves a series of assumptions, i.e., that all conditions are equal away from the farm except for the absence of the farm, and that there have been no changes in background carbon burial nor greater diagenesis since the farm operations started, respectively, which may not be met.

Our findings represent an important step toward quantifying the climate-mitigation cobenefits of seaweed farming. Estimates of carbon burial in sediments below seaweed farms should be included in carbon footprint calculations, which show, even without including this component, that seaweed rank among the farmed food with the lowest carbon footprint (Jones et al. 2022). Our results inform two components required to eventually consider the possibility of issuing blue carbon credits from seaweed farming, (1) additionality, quantified here, as a crude approximation, as excess carbon burial relative to reference sediments, and (2) permanence, as shown from the long-term accumulating of organic carbon documented for the oldest farms in our study. However, permanence requires a commitment to store the organic carbon buried against credits for at least a century. This is much longer than the term of most concessions, requiring further insurances around the requirement of permanence. In addition, robust protocols to verify additional carbon burial below seaweed farms need be developed, as retrospective approaches, such as those used here, cannot be applied to new or recent farms. Moreover, expansion of seaweed farming must be done carefully, based on marine spatial planning informed by hydrological models, to maximize benefits and avoid negative impacts (Duarte et al. 2022a, Berger et al. 2023).

Our estimates are based, however, on average carbon sequestration rates. Our results suggest that operating seaweed farms at high yield in concessions placed in depositional environments, as reflected by the presence of organic-rich sediments, may yield carbon sequestration rates up to four-fold above average. The climate benefits of seaweed farming could be enhanced further by efforts to reduce life cycle emissions and by using the crop retrieved to displace fossil fuels, such as used as biofuel, biochar or goods embodying the seaweed into durable products. Moreover, use of *Asparagopsis* sp. as a feed additive may provide climate benefits by reducing methane emissions by ruminants (e.g. Kinley et al. 2020, De Angelo et al. 2022). Sinking seaweed in the deep sea has been proposed as a path to maximize the climate benefits of seaweed farming (De Angelo et al. 2022). However, this practice may have unintended consequences on the marine ecosystem, while depriving society of the benefits that seaweed crops bring about toward supporting sustainable development goals (Duarte et al. 2022a). Consequently, sinking seaweed in the deep sea is not a recommended option (Ricart et al. 2022).

As the value of carbon credits is projected to raise rapidly over the next years (Ernst & Young 2020), the economic benefits of managing seaweed farms for carbon sequestration will increase in the future and may become, if farmers be compensated for this service, an additional incentive for the expansion of sustainable seaweed farming. However, we submit that, as agricultural production, seaweed farming should be mainly directed to feed humans and/or farmed animals, and should not be treated as a climate geoengineering tool. Rather, enhanced carbon burial in soils below seaweed farming may be considered as a climate mitigation option comparable to that of enhancing organic carbon in agricultural soils (Bossio et al. 2020).

Hence, the (modest) climate mitigation benefits of seaweed farming identified by our results should be considered one more co-benefit to the primary role of seaweed farming in food supply. Documented co-benefits include, in many cases, enhancing biodiversity (Theuerkauf et al. 2022), providing local refugia from ocean acidification and deoxygenation (Xi et al. 2021), along with multiple social benefits, including poverty alleviation and employment (Duarte et al. 2022a). Indeed, the farms in this study collectively employ about 27,000 farmers, corresponding to an average of 1.4 employees per farmed hectare. This provides evidence of the significance of seaweed farming in supporting livelihoods and community resilience in many nations. Indeed, seaweed farming provided food and much needed economic relieve to the farmers in our study when the covid-19 pandemic disrupted other sources of income. For instance, a community heavily dependent on tourism in Bali, Indonesia, switched to seaweed farming to sustain their livelihoods due to the collapse of tourism during the covid-19 pandemic (Pratama and Albasri 2021).

Our results provide the first direct assessment of carbon sequestration below seaweed farms, with an average of 1.87 ton CO_2-eq_ ha^-1^ year^-1^ and a maximum, for the set of farms sampled across the world, of 8.99 ton CO_2-eq_ ha^-1^ year^-1^, which are reduced when using a crude approximation to calculate additionality, to 1.06 and 8.10 ton CO_2-eq_ ha^-1^ year^-1^, respectively. Provided the scalability of sustainable seaweed farming to extend over an ocean space many times larger than the current area (Duarte et al. 2022a), we recommend that knowledge gaps required to assess the climate benefits of seaweed farming across the entire life cycle of the products and other requirements embedded in carbon credits (e.g. Hasselström & Jean-Baptiste, 2022) need be assessed before policies are put in place to incentivate climate change mitigation as a co-benefit of seaweed farming.

## Acknowledgements

This research was funded by ClimateWorks Foundation, the Jeremy and Hannelore Grantham Environmental Trust, the Hindawi Charitable Fund and World Wildlife Fund through grants provided to Oceans 2050’s fiscal sponsor Global Water Challenge, King Abdullah University of Science and Technology (KAUST), and the LIFEWATCH-2019-09-CSIC-13-LWE2021-03-032, funded by the Spanish Ministry of Science and Innovation. Each participating institution provided additional funding for conducting fieldwork, preparing samples for analysis and contributing to the interpretation of results. D.K.-J. was funded by EU H2020 (FutureMARES, contract no. 869300). NNP, AMR and SA were funded in full by World Wildlife ‘Fund and the Bezos Earth Fund. J.W. and X.X. were funded by the Fundamental Research Fund of Zhejiang University (2021XZZX012) and Zhejiang Provincial Natural Science Foundation/Funds for Distinguished Young Scientists (LR22D06003). PIM was supported by an Australian Research Council Discovery Grant (DP200100575). The IAEA is grateful for the support provided to its Marine Environment Laboratories by the Government of the Principality of Monaco. T.K and T.M. were funded by Grants-in-Aid for Scientific Research (KAKENHI, 18H04156 and 18H03354, respectively) from the Japan Society for the Promotion of Science. We thank Inés Sanz Álvarez for her help with analysis, Dr. Chuancheng Fu for help with figures, Dr. Kenta Watanabe, Dr. Hirotada Moki, Ms. Toko Tanaya, Dr. Naoko Morimoto for help with sample processing for the Japanese farms. and the many farmers and volunteers that helped with sampling and sample processing.

## Methods

### Sampling

A network of 20 seaweed farms worldwide was assembled, usually including a local research team operating under standardized protocols. The current species farmed and the recent yields were provided by the farm operators for each farm, along with the time since farming operations started. The oldest and largest farms available were selected for sampling, although relatively recent and small farms in Europe and North America, where seaweed farming is a recent development, were included. For each of the selected farms we defined a “farm” site, representing a site that is likely to receive continuous inputs of any materials detached from the farm, and a “reference” site, representing a site that is similar in depth and overall environmental conditions (e.g., exposure) as the farm site, but sufficiently away as to be unlikely that materials from the farm may settle there. The “farm” site was located below the center of the farm area, where “edge” effects will be minimal. Where the farm area shifted over time, the “farm” site represented the area that best represent the center across years (i.e., consistently covered by the farm). Where significant directional current occurred and depth was significant (> 10 m), the “farm” site was displaced relative to the center of the farm along the direction of the prevailing current, as settling materials will be advected along that direction, rather than falling vertically. Both sites were also located away from known disturbances to the sediment, that can mix or rework the sediments, such as anchoring systems.

A total of 3 sediment cores, ranging in length from 60 cm to 1 m and 6 to 10 cm in dimater, were collected at each of the farm sites and 2 additional cores at the reference sites, when possible just before the crop was harvested. Cores were collected by SCUBA divers at most farm, but for deep farms the use of a vibracorer operated from a vessel equipped with an A frame was necessary. As sampling was set to be initiated in March 2020, covid-19 restrictions led to significant constraints in accessing the sites for research teams due to confinement and restriction measures under covid19, which were severe in some of the participating farms, such as those in China and Japan. Hence, sediment sampling elapsed between March 2020 and June 2022, depending on farms. The farm in the US was sampled twice, and the average values from the two sampling events are provided.

The cores retrieved were sliced at 0.5 cm intervals for the top 10 cm, 1 cm intervals from 10 cm to 20 cm, and 2 cm intervals down to the base of the core. The wet weight of the slices was determined, and they were then dried at 60-70 °C until constant weight. Dried samples were then shipped to the laboratories for analyses. Organic carbon was determined at the Andalusian Institute of Earth Sciences (Granada, Spain), and ^210^Pb concentrations were determined, at Edith Cowan University (Perth, Australia), and the Marine Environment Laboratories of the UN International Atomic Energy Agency in Monaco.

### Chemical analyses

Up[on reception in the laboratories, sediment samples were dried again at 70 °C to remove any water absorbed during the shipment and subsequently pulverized/homogenized. An aliquot was treated with HCl (1:1) to remove carbonates. The residual sample was washed (5 times) with MiliQ water, dried and homogenized again, to analyze the percentage of organic carbon. Acid treatment and washing may have removed some soluble organic carbon, such as soluble polysaccharides, possibly leading to underestimating organic carbon (Komada et al. 2008). However, the risk in reducing acid treatment would have been to overestimate organic carbon by failing to dissolved carbonates, that dominate sediment material in tropical sediments. Samples were accurately weighed into tin capsules and analyzed in triplicate to calculate the carbon content by means of a Carlo Elba NC1500 (Milan, Italy) elemental analyzer coupled with a Delta Plus XP (Thermo-Finnigan, Bremen, Germany) mass spectrometer (EA-IRMS). Commercial CO_2_ was used as the internal standard for the carbon analyses. Mass area 44, 45 and 46 of CO_2_ were used to calculate the carbon content.

The concentrations of ^210^Pb along each core were determined by alpha spectrometry through the measurement of its granddaughter ^210^Po, assuming radioactive equilibrium between both radionuclides (Sanchez-Cabeza et al., 1998). Briefly, 300 mg of each sample were spiked with a precisely known amount ^209^Po and microwave digested with a mixture of concentrated HNO3 and HF. Boric acid was then added to complex fluorides. The resulting solutions were evaporated and diluted to 100 mL 1M HCl and Po isotopes were plated onto pure silver disks during 8 hours at 70 °C while stirring. Polonium emissions were measured by alpha spectrometry using ULTRA Ion implanted silicon charged particle detectors (ORTEC). The concentrations of supported ^210^Pb were determined in several evenly distributed samples along each core by measuring the concentrations of ^226^Ra by gamma spectrometry via its decay products ^214^Pb and ^214^Bi (in equilibrium) using well-type, high purity germanium detectors (CANBERRA). Analyses of replicate samples and reference materials (IAEA-447) were carried out systematically to ensure the accuracy and the precision of the results for both alpha and gamma measurements. The excess ^210^Pb concentration profiles were obtained by subtracting the supported ^210^Pb from the total ^210^Pb for each core. Mean sedimentation rates were obtained by modeling the excess ^210^Pb concentration profiles along the accumulated mass using the CF:CS model (Krishnaswamy et al. 1971, Arias-Ortiz et al., 2018) where possible (i.e. where the excess ^210^Pb concentration profile presented a decreasing trend with depth). Where a mixed layer existed in the upper layer of the sediment, sedimentation rates calculated in the underlaying layer were assumed to apply to the overlaying mixed layer (Arias-Ortiz et al., 2018).

### Calculations

Organic carbon (C_org_) concentrations were combined with bulk density, calculated as the ration of the dry weight and the fresh volume of the sediment sections, to calculate carbon density and C_org_ stocks in the sediments. For each core where the age model could be calculated, the C_org_ burial rate since the beginning of activity of the farm was estimated using the mean sedimentation rate and the weighted average concentration of C_org_ from the surface and down to the depth corresponding to the time elapsed, as calculated from the age model. Similarly, the C_org_ burial rate during the equivalent amount of time before the farm was operative was estimated using the mean sedimentation rate and the weighted average concentration of C_org_ at layers below those estimated to correspond to the onset of farming operations of each sediment core collected below the seaweed farm. Then, where possible, we could compare the C_org_ burial rates during and before operation of the farm as well with the C_org_ burial rates outside the farms (i.e., determined from samples collected in reference sites). The thickness and carbon stock deposited during farm operations were extrapolated linearly from the sediment layer analyzed (Table 2) when this did not include the entire operation period of the farm. Examination of sediment properties (e.g., bulk density, organic concentration) showed that “reference” values, both those sampled beyond the farm and those retrieved from sediment layers deposited before farm operations were initiated, were imperfect proxies for the carbon burial in sediments in the absence of farming due to spatial and temporal variability. In some farms (Dongtou, China, and the farms sampled in Canada), the sediments in reference cores were of a different nature than those in the farm before the onset of farm operations, so the best reference was provided by the data from sediments deposited before the onset of farm operations, while sediments beyond the farm were used as best proxies of “reference” sediments for all other farms. This was not possible for very old farms or those supporting very high sedimentation rates, where the sediment horizons prior to the onset of farming activities were not reached. The farm in the US was sampled twice, and values reported represent average from the two sampling events.

## Data Availability

Data are provided in Tables 1 and 2. ^210^Pb concentrations will be made available in an open access data repository upon acceptance of the paper, and will be included in reference section with the appropriate DOI.

